# Multiple clines within an ecotype of the yellow monkeyflower, *Mimulus guttatus*

**DOI:** 10.1101/2023.07.24.550335

**Authors:** Thomas Zambiasi, David B. Lowry

## Abstract

**Premise:** A key goal of evolutionary biologists is to understand how and why genetic variation is partitioned within species. In the yellow monkeyflower, *Mimulus guttatus* (syn. *Erythranthe guttata*), coastal perennial populations collectively constitute a single genetically and morphologically differentiated ecotype compared to inland populations of *M. guttatus*. While the distinctiveness of the coastal ecotype has now been well documented, there is also variation in environmental factors across the range of the coastal ecotype that could drive differentiation among its component populations in a more continuous way.

**Methods:** Based on previous observations of a potential cline within this ecotype, we quantified plant height across coastal perennial accessions from 74 total populations in a greenhouse common garden experiment. To evaluate possible environmental factors driving the relationship between trait variation and latitude, we regressed height against multiple climatic factors, including temperature, precipitation, and coastal wind speeds.

**Results:** Multiple traits were correlated with latitude of origin, but none more than plant height. Plant height was negatively correlated with latitude and plants directly exposed to the open ocean were shorter than those that were more protected from onshore coastal winds. Further analyses revealed that height was correlated with climatic factors (precipitation, temperature, and windspeeds) that were autocorrelated with latitude. We hypothesize that one or more of these climatic factors drove the evolution of latitudinal clinal variation within the coastal ecotype.

**Conclusion:** Overall, our study illustrates the complexity of how the distribution of environmental variation can simultaneously drive the evolution of distinct ecotypes as well as continuous clines within those ecotypes.

## INTRODUCTION

Over a century ago, Turesson (1922a,b) recognized that adaptive phenotypic variation in plants was often highly divergent in correlated sets of traits when comparing groups of populations derived from different habitats. Turesson referred to these groups of phenotypically similar, locally-adapted populations as ecotypes (Turesson 1922a,b; Gregor et al. 1936). Several researchers following Turesson found the concept of ecotypes useful for describing the distribution of adaptive genetic variation within species (Gregor et al. 1936; Clausen et al. 1948; Clausen 1951; Böcher 1967; Heywood 2011). Further, these researchers viewed partially reproductively isolated ecotypes as an intermediate stage in the process of speciation and a way to directly link natural selection to the formation of new species (reviewed in Lowry 2012).

However, other researchers doubted the validity of Turesson’s ecotype concept and argued that clines (Huxley 1938) best represented how adaptive genetic variation is partitioned across the geographic distribution of a species’ range (Langlet 1963; 1971; Stebbins 1980).

While many contemporary biologists recognize the utility of concepts like ecotypes and/or clines to describe the distribution of adaptive phenotypic variation within species, there is still a lack of clarity of the full extent to which phenotypic variation is distributed across the complexity of the natural landscape (Lowry et al. 2014; Briggs and Walters 2016). In particular, questions remain about the extent to which geographically widespread ecotypes themselves may harbor clines in phenotypic variation. This question was of major importance to the biosystematists like Clausen, Keck, Hiesey, (Clausen et al. 1948) and their contemporaries (Gregor 1930, 1938), but has since received limited attention.

In this study, we examined the distribution of phenotypic variation within a widespread ecotype of the yellow monkeyflower, *Mimulus guttatus* (syn. *Erythrante guttata*; Barker et al. 2012)*. M. guttatus* exhibits high levels of genetic and phenotypic variation across its extensive range spanning western North America, which makes it a particularly useful system for studying how local adaptation shapes the distribution of adaptive genetic variation across space (Vickery 1978; Lowry et al. 2008). Coastal perennial populations of *M. guttatus*, occurring from southern California to southern Washington State, collectively comprise a distinct ecotype based on population genetic data (Lowry et al. 2008; Twyford and Friedman 2015). This ecotype has been designated as a variety (*M. guttatus var. grandis* Greene) or species (*M. grandis* syn. *E. grandis*; Barker et al. 2012, Nesom 2012, 2014) by taxonomists. Despite these taxonomic designations due to their morphological distinctiveness, the coastal perennial ecotype is highly interfertile with the inland annual ecotype of *M. guttatus* (Lowry and Willis 2010). Further, there were only four fixed genetic differences between the coastal perennial and inland annual populations out of 29,693,578 SNPs surveyed in a population genetic study of this system (Gould et al. 2017).

Based on these results, we believe that it is best to think of the coastal perennial populations as an intraspecific ecotype within the *Mimulus guttatus* species complex, which is the result of non-fixed differences in allele frequencies at many loci across the genome (Lowry and Willis 2010; Twyford and Friedman 2015; Gould et al. 2017).

Several previous studies of local adaptation in *M. guttatus* have found striking genetically-based phenotypic differences between the coastal perennial ecotype and nearby inland populations in the western United States (Vickery 1952; Hall et al. 2006, 2010; Lowry et al. 2008; Oneal et al. 2014). Inland *M. guttatus* populations of the coastal mountain ranges of western North America primarily have an annual life-history, especially those that occur in inland habitats that completely dry out during summer months (Vickery 1952; Clausen and Heisey 1958; Wu et al. 2008; Lowry et al. 2008). While inland populations in the coast ranges are annuals, perennial populations do occur with some frequency in areas that maintain soil water through the summer months (van Kleunen 2007; Oneal et al. 2014; Twyford & Friedman 2015). The occurrence of inland perennial populations increases further inland, especially at higher elevations in the Sierra Nevada and Cascade Mountain ranges (Coughlin et al. 2021). The level of morphological differentiation between coastal perennial and inland annual ecotypes is immense (Lowry & Willis 2008). Further, coastal perennial populations are separated genetically from inland populations, both annual and perennial, in previously published principal component plots (Fig. 3 of Lowry 2012 and Fig. S2 of Twyford & Friedman 2015). In contrast to the inland annuals, the coastal perennial ecotype occurs in coastal habitats that are regularly damp and foggy during summer months (Vickery 1952; Clausen and Heisey 1958; Hitchcock and Cronquist 1973; Hall and Willis 2006; Lowry et al. 2008). In greenhouse common garden experiments, accessions derived from coastal habitats had thicker stems, many more prostrate stoloniferous branches, later flowering times, and produced far more herbivore defensive compounds than nearby inland annual populations (Hall et al. 2006, 2010; Lowry et al. 2008; Lowry and Willis 2010; Oneal et al. 2014; Lowry et al. 2019).

Much of the trait differences between the inland annual and coastal perennial ecotypes contribute to local adaptation (Lowry et al. 2008, 2019; Lowry & Willis 2010; Popovic & Lowry 2019) and are caused by a few large effect loci, including a large chromosomal inversion polymorphism on chromosome 8 (Lowry and Willis 2010; Oneal et al. 2014; Twyford & Friedman 2015; Lowry et al. 2019). Inland perennial populations of *M. guttatus* consistently have the same orientation of the chromosomal inversion as coastal perennial populations (Lowry & Willis 2010). Indeed, one of the few regions of the genome that is highly differentiated between inland annual and inland perennial populations is the chromosome 8 inversion (Oneal et al. 2014; Twyford & Friedman 2015). In this study, we focus on phenotypic variation only among coastal perennial populations, which we assume all share the same orientation chromosome 8 orientation based on previous studies (Lowry and Willis 2008; Oneal et al. 2014; Twyford & Friedman 2015; Gould et al. 2017; Kollar et al. 2023).

While there is both phenotypic and genetic divergence between ecotypes of *M. guttatus*, it is unclear to what extent genetic variation is present within individual ecotypes – either due to adaptation to widespread environmental gradients or idiosyncratic with regard to geographic space. However, a previous pilot study suggested the possibility that there may be a latitudinal cline in plant height within the coastal ecotype (Lowry 2010). Since that initial study, dozens of additional coastal *M. guttatus* populations have been collected (Fig. 1), which provides an opportunity for a more comprehensive survey of variation over the entire geographic range of the coastal perennial ecotype.

**Figure 1.**
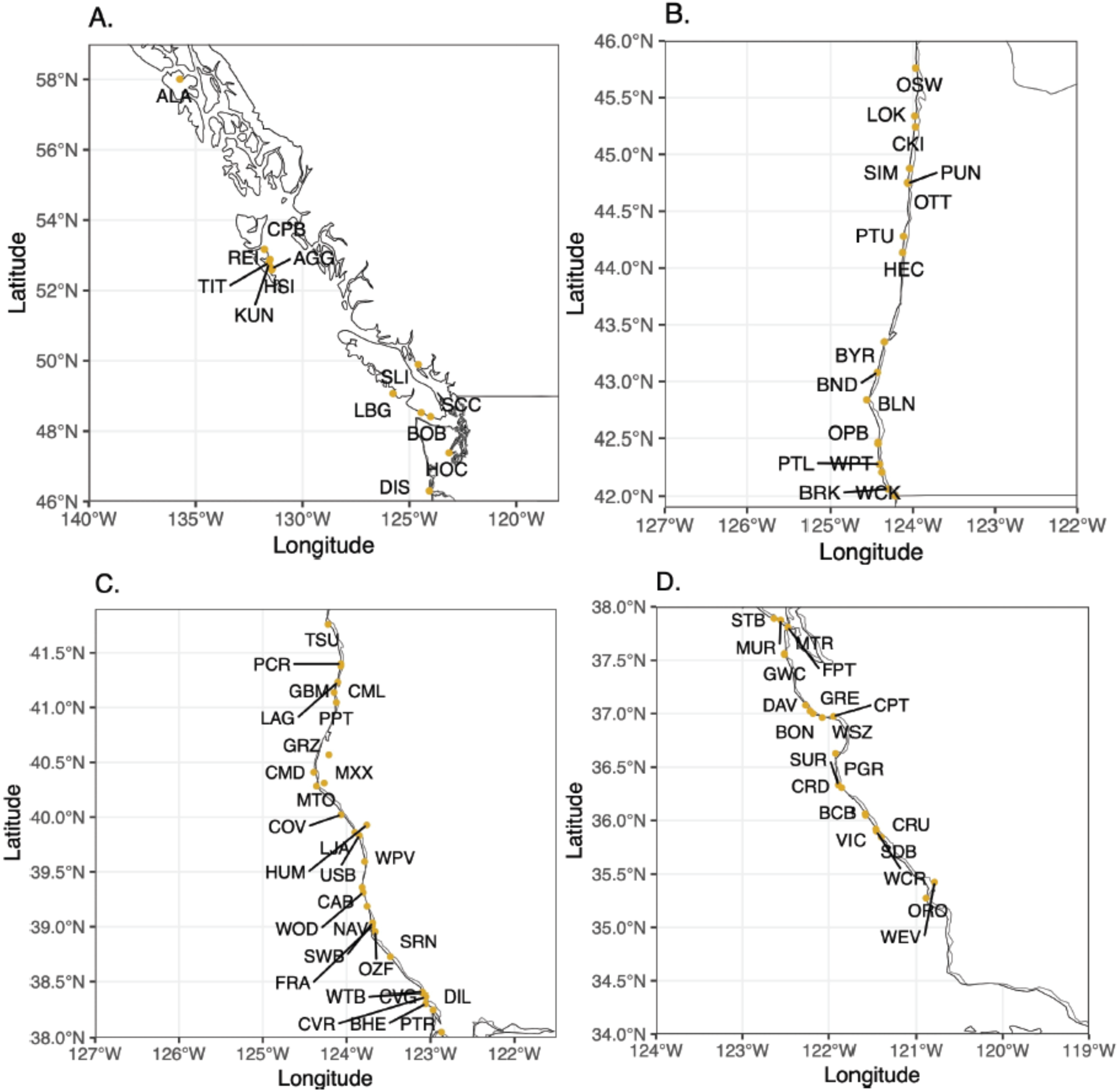
Population locations for coastal perennial *M. guttatus* in a) Washington, British Columbia, and Alaska; b) Oregon; c) Northern California; and d) Southern California. These maps include populations grown in both the main and pilot studies.

In this study, we tested the hypothesis that there is a latitudinal cline in phenotypic variation within the coastal ecotype. We simultaneously evaluated the role of exposure of the coastal perennial populations to the open ocean on trait variation. To establish potential sources of selection, we examined whether there is a relationship between the phenotypes of the accessions and the climates from where they were collected in nature. Through these efforts, we attempted to understand whether clinal trait variation occurred within the distinctive coastal perennial ecotype.

## METHODS

### Germplasm for experiments

The seeds used in these experiments originated from natural population collections made between 2004 and 2016. Seeds were either collected directly from plants in the field or collected from plants that had been brought from the field back to the greenhouses at Duke University (Durham, North Carolina, USA) or Michigan State University (East Lansing, Michigan, USA). All seeds used for the experiment were grown for at least one generation in the greenhouse prior to the study. For each population, we collected location description information, habitat description, and GPS coordinates. Photos of the plants in the field for each population have been posted on iNaturalist and vouchers for a subset of the populations were deposited at the California Academy of Sciences. Permits were obtained for collections on public lands, where permission was required. All maternal families used in this study are currently stored at Michigan State University in airtight containers with silica gel as a desiccant.

### Greenhouse Experiment

To determine whether there was clinal variation in *M. guttatus* traits, we conducted an experiment in a controlled greenhouse environment. We conducted the greenhouse experiment in the Plant Ecology Field Lab greenhouse at Kellogg Biological Station (KBS, Hickory Corners, MI, USA). Seeds from coastal perennial *M. guttatus* (*N* = 423 individuals, 418 seed families, 74 populations) were planted with SUREMIX (Michigan Grower Products, Inc., Galesburg, Michigan, USA) in 3.5 sq. in. pots without prior cold stratification. We included a variable number of seed families per population (Range = 1-15, Mean = 5.72 families) and all families but five were represented by a single individual (BON-2, BON-4, CPT-1, CPT-5, and CRU-1 each had two individuals measured). More information on the populations used in this experiment, plus the coordinates and maps of the original population locations can be found in Figure 1 and Table S1 in the supplementary material (Appendix S1; see Supplemental Data with this article). All germinating seedlings except the centermost plant were removed from the pot and remaining individuals were watered every other day throughout the experiment. Germination and flowering dates were recorded to determine flowering time; height of plant at first flower (cm), width of the second set of true leaves (cm), corolla length (mm), and corolla width (mm) were measured at the time of first flowering.

### Climate Data

To investigate potential environmental drivers of adaptive phenotypic variation, we extracted climatic data for each population’s collection location of origin from the WorldClim Bioclimatic variables database (Fick and Hijmans 2017). This data was imported and compiled using R (v4.2.1, R Core Team 2022) using the *stats* (v4.2.1), *nlme* (v3.1-157)*, raster* (v3.6-26), and *sp* (v2.1-3) packages.

In addition to the Bioclim data, we extracted oceanic wind speed data from along the US Pacific Coast using the National Renewable Energy Laboratory’s Wind Prospector website, which featured interactive online maps of wind speeds surveyed by NREL (King et al. 2014; Draxl et al. 2015a,b). We used Pacific Monthly Offshore Wind Speed map layers to collect wind speeds for each month from the site nearest to shore (8 km offshore at the closest; Draxl et al. 2015b) at the latitudes of each of the coastal perennial populations. The wind speeds found through the NREL Wind Prospector were the averages for each 1.2 km X 1.2 km offshore site and were modeled at 100 m (Draxl et al. 2017). Due to unavailability of wind data outside of the contiguous US, populations found in British Columbia and Alaska (denoted by population codes ALA and LBG) were not included in climate data analyses. Monthly wind speed averages accessible via NREL were based on observations and statistical analyses over the course of seven years (King et al. 2014; Draxl et al. 2015a,b), which we used to calculate quarterly and annual wind speeds for each population collection location. We then matched these climate variables with their corresponding coordinates and population codes for further analysis.

### Statistical Analysis

Prior to data analyses, we calculated mean trait values for each population to avoid pseudoreplication in our analyses. Our analyses included a binary factor to identify whether each population was exposed to (1) or protected (0) from the open ocean. Exposed populations were located on the cliffs, headlands, and seeps just adjacent to the open Pacific Ocean. In contrast, protected populations were not directly exposed to the open ocean by being located in inland roadside ditches and lagoons or on the coastlines of Washington, British Columbia, and Alaska in locations where they were protected from the open ocean by other land masses. To test whether there was a relationship between plant traits and the latitude of origin (location from which each population accession was collected), we modeled trait values against latitude with the binary predictor for ocean exposure via linear regression in R (v4.2.1, R Core Team 2022) using the *stats* (v4.2.1) package, evaluated predictor significance via ANOVA (*car* package v3.1-2), and visualized our results in *ggplot2* (v3.5.0). We modeled both additive and interactive relationships between the predictors for each trait, but in each case found that the additive model was a better descriptor of the data per AIC and ANOVA model comparison (both in the *stats* package v4.2.1).

Although we predicted that latitude would correlate with plant height across space, we wanted to evaluate which climatic variables correlated with latitude might ultimately be driving observed clinal patterns. Climate variables are often highly correlated with each other, so we conducted a principal components analysis (PCA) to reduce the dimensionality of the data prior to further analyses. We extracted climate data, described above, for each population and performed the PCA in R (v4.2.1) using the *factoextra* (v1.0.7) and *stats* (v4.2.1) packages. We evaluated all principal components (PCs) with an eigenvalue greater than 1, following the Kaiser criterion (Kaiser 1960). We then conducted regression analyses of population mean plant height against each of these PCs individually and with all as factors in the same model, again including the exposure factor and using linear models in the *stats* (v4.2.1) package. We also evaluated models with interactions and polynomial fits and selected the best descriptors for the data via AIC and ANOVA model comparison. For all regression analyses, we evaluated residuals with the *DHARMa* (v.0.4.6) package and confirmed via visual inspection that residuals were normally distributed for all models. All data organization, analysis, and visualization was conducted in R Studio (2023.12.0+369 “Ocean Storm” Release, Posit Team 2023).

## RESULTS

### Trait correlations with latitude

We found a significant negative relationship between latitude of population origin and plant height within the coastal perennial ecotype in our greenhouse experiment (*R^2^* = 0.43, *F*_2,71_ = 26.83, *P* < 0.001; Figs. 2, 3). Based on our linear modeling, we found that every degree latitude further north translated to a roughly 1 cm decrease in plant height under greenhouse conditions. We also found a significant difference in the height-latitude relationship depending on whether populations were highly exposed to the wind and salt spray from the open Pacific Ocean. In general, protected populations were taller than those exposed to the open ocean (Table 1, Fig. 2). In addition to the strong relationship with height, we found significant, but much weaker relationships between population latitude and width of the second set of true leaves (*R^2^* = 0.12, *F*_2,71_ = 4.83, *P* = 0.011), corolla width (*R^2^* = 0.12, *F*_2,63_ = 4.38, *P* = 0.017), and corolla length (*R^2^* = 0.17, *F*_2,63_ = 6.51, *P* = 0.003). Leaf width generally decreased with latitude, as did corolla width and length; none of these models included a significant term for ocean exposure (Table 1). The relationship of latitude to flowering time (*R^2^* = 0.03, *F*_2,71_ = 0.97, *P* = 0.383) was not significant. Because the correlation between plant height and latitude was so much stronger than for the other traits, we focused the rest of our analyses on the possible causes of this correlation.

**Figure 2.**
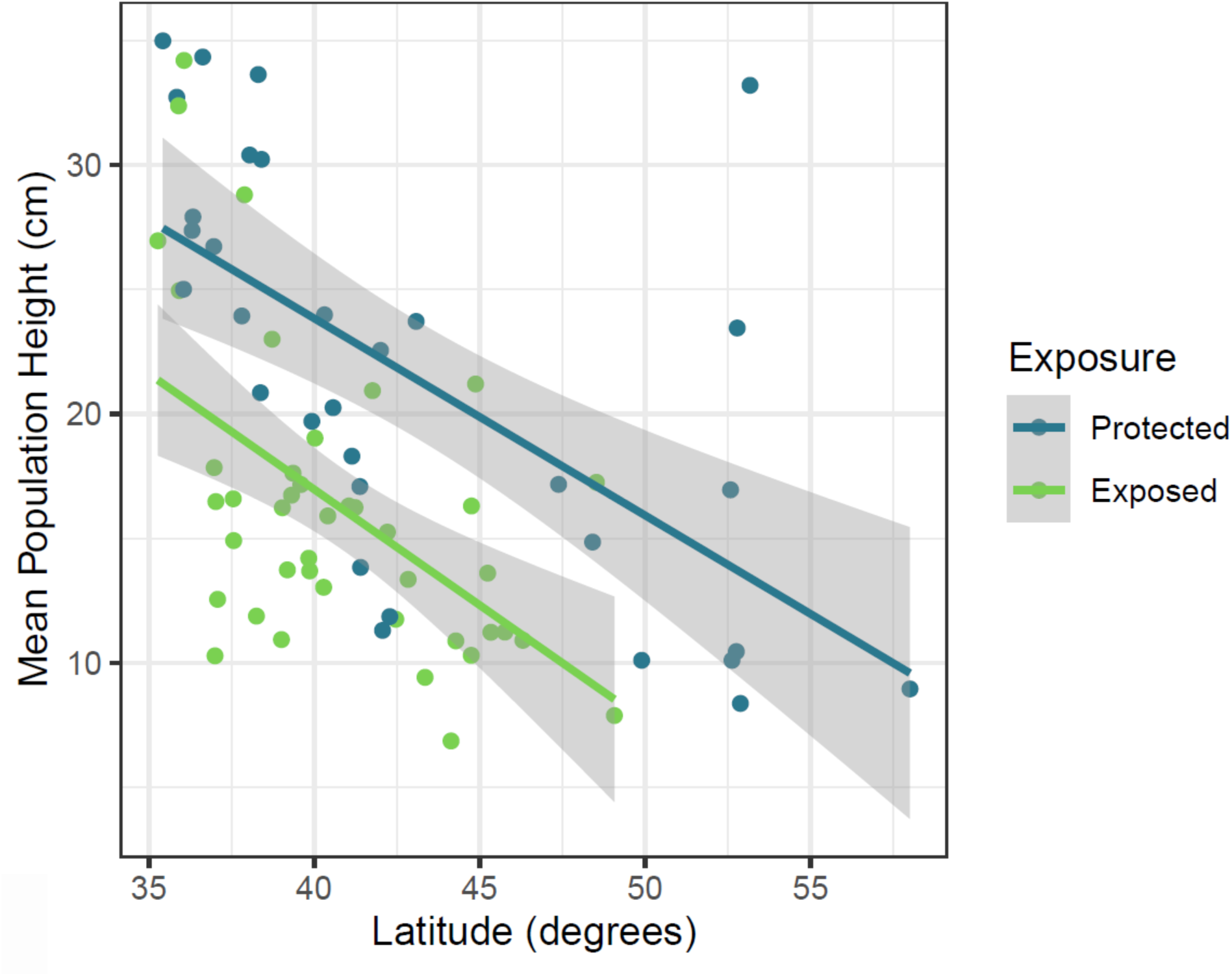
Mean *M. guttatus* population height decreases significantly with latitude (*P* < 0.001), and exposed populations have shorter average heights than protected populations (*P* < 0.001); both population groups express the same relationship between height and latitude of origin. Relationship of latitude of origin (in degrees) for *M. guttatus* populations and mean individual heights (cm) of these populations was determined by linear regression.

**Figure 3.**
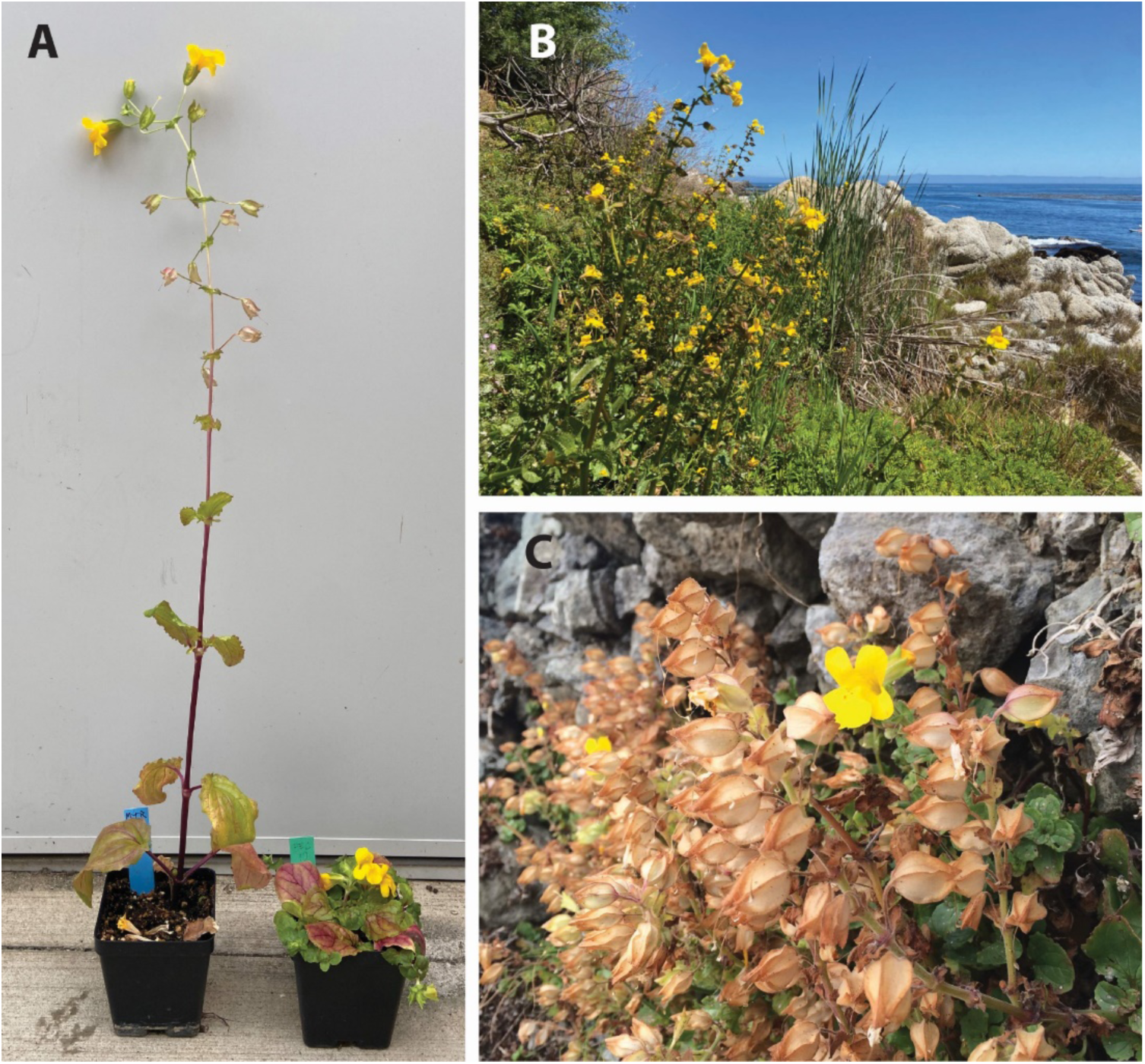
Variation in plant height within the coastal perennial ecotype in *Mimulus guttatus.* A) Divergence in height between plants from the Central California coast (left) and Oregon (right) when grown in a greenhouse common garden experiment. B) Tall plants growing in a protected area of the Monterey Bay along the Central California coast reach up to 2 meters in height. C) Plants exposed to the open ocean in Oregon are much more compact in growth form.

**Table 1.** ANOVA and linear regression summaries for *M. guttatus* traits in the main greenhouse. . Asterisks denote significance at p<0.05 (*), p<0.01 (**), and p<0.001 (***).

| Trait | Parameter Summaries |  |  |  |  |  | Full Model Summaries |  |  |  |
| --- | --- | --- | --- | --- | --- | --- | --- | --- | --- | --- |
|  | Predictor | Estimate | Sum Sq. | df | F | P | Multiple R<sup>2</sup> | df | F | P |
| Flowering Time (days) | Latitude | -0.0633 | 7.6 | 1,71 | 0.120 | 0.7301 | 0.0267 | 2,71 | 0.974 | 0.3825 |
|  | Exposure | 2.3836 | 98.4 | 1,71 | 1.551 | 0.2171 |  |  |  |  |
| 2nd Leaf Width (cm) | Latitude | -0.1633 | 50.56 | 1,71 | 9.661 | 0.0027** | 0.1198 | 2,71 | 4.83 | 0.0108* |
|  | Exposure | -0.3736 | 2.42 | 1,71 | 0.462 | 0.4990 |  |  |  |  |
| Corolla Width (mm) | Latitude | -0.2036 | 71.82 | 1,63 | 5.636 | 0.0207* | 0.1221 | 2,63 | 4.383 | 0.0165* |
|  | Exposure | 1.0034 | 15.13 | 1,63 | 1.187 | 0.2800 |  |  |  |  |
| Corolla Length (mm) | Latitude | -0.2106 | 76.84 | 1,63 | 7.008 | 0.0102* | 0.1713 | 2,63 | 6.511 | 0.0027** |
|  | Exposure | 1.4334 | 30.87 | 1,63 | 2.816 | 0.0983 |  |  |  |  |
| Height (cm) | Latitude | -0.8297 | 1305.94 | 1,71 | 39.017 | 2.738e-08*** | 0.4304 | 2,71 | 26.83 | 2.099e-09*** |
|  | Exposure | -7.0716 | 865.79 | 1,71 | 25.867 | 2.863e-06*** |  |  |  |  |

### The relationship of climatic factors and plant height

In our greenhouse experiment, we found that climate PC1 through PC3 all had eigenvalues greater than 1. In total, those three PCs accounted for 89.99% of cumulative variance in climate factors. PC1 explained the most variation at 56.39%, followed by PC2 with 22.81%, and PC3 with 10.79%. PC1 was significantly negatively correlated with plant height in a linear regression model (*PC1 estimate* = -1.09, *exposure estimate* = -5.69, *Multiple R²* = 0.52, *F*_2,62_ = 33.05, *P* < 0.001). Due to the nonlinear relationship between wind speeds and latitude, we also assessed nonlinear regression models and found that the relationship between PC1 and plant height was best explained by a quadratic regression model as determined via AIC (*Multiple R^2^* = 0.57, *F_3,61_* = 27.23, *P* < 0.001; *AIC linear* = 402.52, *AIC quadratic* = 396.46). The quadratic regression yielded the same qualitative relationships between mean population height and the climate factors as the linear model; height decreased with increasing values of PC1 (Fig. 4). We also evaluated regression models of the remaining PCs against plant height. While each showed a significant relationship with height, PCs 2 and 3 only included significant terms for ocean exposure (Table 2). However, our model with the PC1 square term and ocean exposure as predictors was a better descriptor of the data based on AIC than models with PC2 and PC3 (Table 2). We also constructed a linear model with PCs 1-3 and exposure as predictors to compare its fit with our quadratic PC1 model, but yet again found PC1 to best describe the data via AIC (*AIC PC1* = 396.46, *AIC All PCs* = 396.82). Following our tests for model fit, we focused on PC1’s loadings for the remainder of our interpretations.

**Figure 4.**
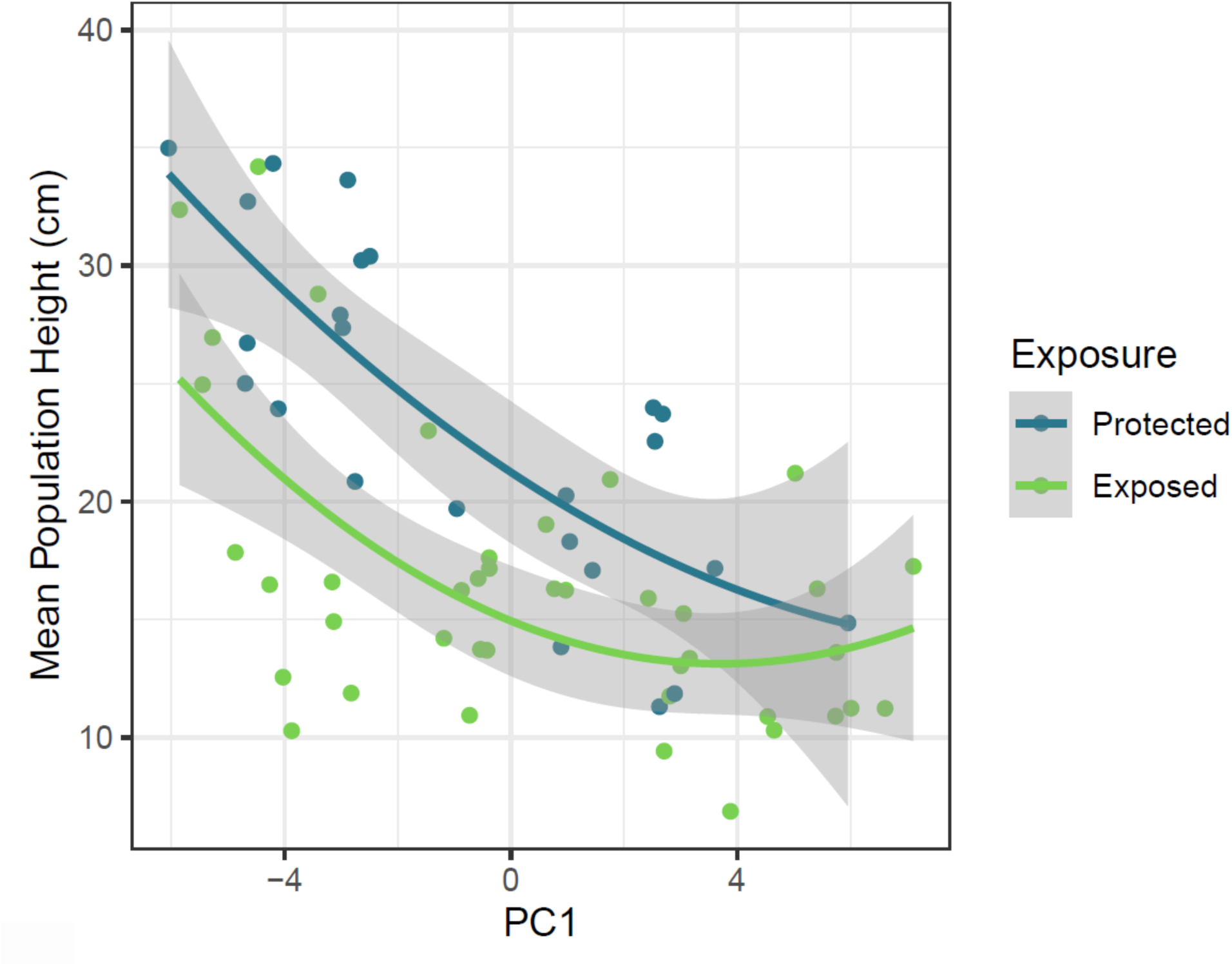
Mean *M. guttatus* population height decreased significantly and non-linearly with increasing values of climate PC1 in the main experiment (*P* < 0.001); exposed populations were also significantly shorter than protected populations but maintained the same relationship to PC1 (*P* < 0.001). The figure displays the quadratic relationship between climate PC1 and plant height, which was the best fitting model as determined by AIC.

**Table 2.** ANOVA, linear regression, and AIC summaries for models evaluating the relationship of selected principal components to *M. guttatus* height in the main greenhouse. Asterisks denote significance at p<0.5 (*), p<0.01 (**), and p<0.001 (***). Lowest AIC is bolded.

| Principal Component | Parameter Summaries |  |  |  |  |  | Full Model Summaries |  |  |  | AIC |
| --- | --- | --- | --- | --- | --- | --- | --- | --- | --- | --- | --- |
|  | Predictor | Estimate | Sum Sq. | df | F | P | Multiple R<sup>2</sup> | df | F | P |  |
| PC1 | (Dim. 1)<sup>2</sup> | 13.923 | 1193.90 | 2,61 | 25.044 | 1.147e-08*** | 0.5725 | 3,61 | 27.23 | 2.679e-11*** | **396.462** |
|  | Exposure | -6.048 | 533.47 | 1,61 | 22.381 | 1.368e-05*** |  |  |  |  |  |
| PC2 | Dim. 2 | 0.0520 | 0.94 | 1,62 | 0.022 | 0.883 | 0.2217 | 2,62 | 8.831 | 4.219e-04*** | 433.403 |
|  | Exposure | -7.0733 | 752.57 | 1,62 | 17.628 | 8.736e-05*** |  |  |  |  |  |
| PC3 | Dim. 3 | 0.0663 | 0.71 | 1,62 | 0.017 | 0.898 | 0.2217 | 2,62 | 8.828 | 4.23e-4*** | 433.408 |
|  | Exposure | -7.0856 | 743.63 | 1,62 | 17.417 | 9.524e-05*** |  |  |  |  |  |

We evaluated which climatic factors loaded onto PC1. Heavily loaded climate variables included Annual Precipitation, Average Annual Temperature, Precipitation Seasonality, Precipitation of the Warmest Quarter, Precipitation of the Wettest Quarter, Precipitation of the Coldest Quarter, Precipitation of the Wettest Month, and Precipitation of the Driest Quarter; Average Wind Speeds in Quarter 1 (January, February, March) and Quarter 4 (October, November, December) were not as heavily loaded, but were still represented on PC1 (Table S2). Higher values of PC1 indicated lower Average Annual Temperature and Precipitation Seasonality, but greater Annual Precipitation, Precipitation of the Warmest Quarter, Precipitation of the Wettest Quarter, Precipitation of the Coldest Quarter, Precipitation of the Wettest Month, Precipitation of the Driest Quarter, and Average Wind Speeds in Quarter 1 and Quarter 4 (Table S2). Therefore, any population at a location corresponding to a high value of PC1 experienced lower temperatures, higher amounts and greater consistency of precipitation, and faster wind speeds than populations at sites with a low PC1 value.

### Latitude strongly predicts climate for coastal populations

To assess how well latitude predicts climate at the sites of our study populations, we regressed the extracted principal components from our greenhouse experiment against latitude. Our analysis confirmed that many of the climate variables involved were highly correlated with latitude. PC1 was the only principal component significantly associated with latitude and the two variables had a positive relationship (*slope* = 0.89, *Multiple R²* = 0.92, *F*_1,63_ = 704.9, *P* < 0.001). To determine how well variation in individual climatic factors was predicted by latitude, we regressed all the climate factors that were highly loaded on PC1 individually against latitude (Table 3).

**Table 3.** Regressions of climate factors that were highly-loaded on PC1 against latitude.

| *Climate Factors* | Estimate | F | Multiple R<sup>2</sup> | P |
| --- | --- | --- | --- | --- |
| Annual Precipitation | 0.0495 | 231 | 0.7857 | < 2.2e-16 |
| Coldest Quarter Precipitation | 0.1116 | 136.4 | 0.684 | < 2.2e-16 |
| Driest Quarter Precipitation | 0.6088 | 405.3 | 0.8655 | < 2.2e-16 |
| Mean Annual Temperature | -2.5849 | 350.2 | 0.8475 | < 2.2e-16 |
| Mean Wind Speed, 1st Quarter | 2.4760 | 53.32 | 0.4584 | 5.94E-10 |
| Mean Wind Speed, 4th Quarter | 1.8632 | 64 | 0.5039 | 3.57E-11 |
| Precipitation Seasonality | -0.3049 | 549.7 | 0.8972 | < 2.2e-16 |
| Warmest Quarter Precipitation | 0.6092 | 483.8 | 0.8848 | < 2.2e-16 |
| Wettest Month Precipitation | 0.3099 | 137.1 | 0.6852 | < 2.2e-16 |
| Wettest Quarter Precipitation | 0.1054 | 159.6 | 0.7169 | < 2.2e-16 |

Note: F-statistic degrees of freedom were all 1,63

### Patterns of wind speed along the west coast of North America

Plants are well known to respond to wind through changes in growth form (Grace 1977; Gardiner et al. 2016; Telewski 2021). This, combined with the observation of wind speed loading onto the first PC of climate, led us to evaluate the patterns of wind speed variation more carefully across the populations that we studied (Fig. 5). We found that, in general, annual wind speeds increase with latitude (*slope* = 0.07, *Multiple R²* = 0.06, *F*_1,76_ = 4.465, *P* = 0.038).

**Figure 5.**
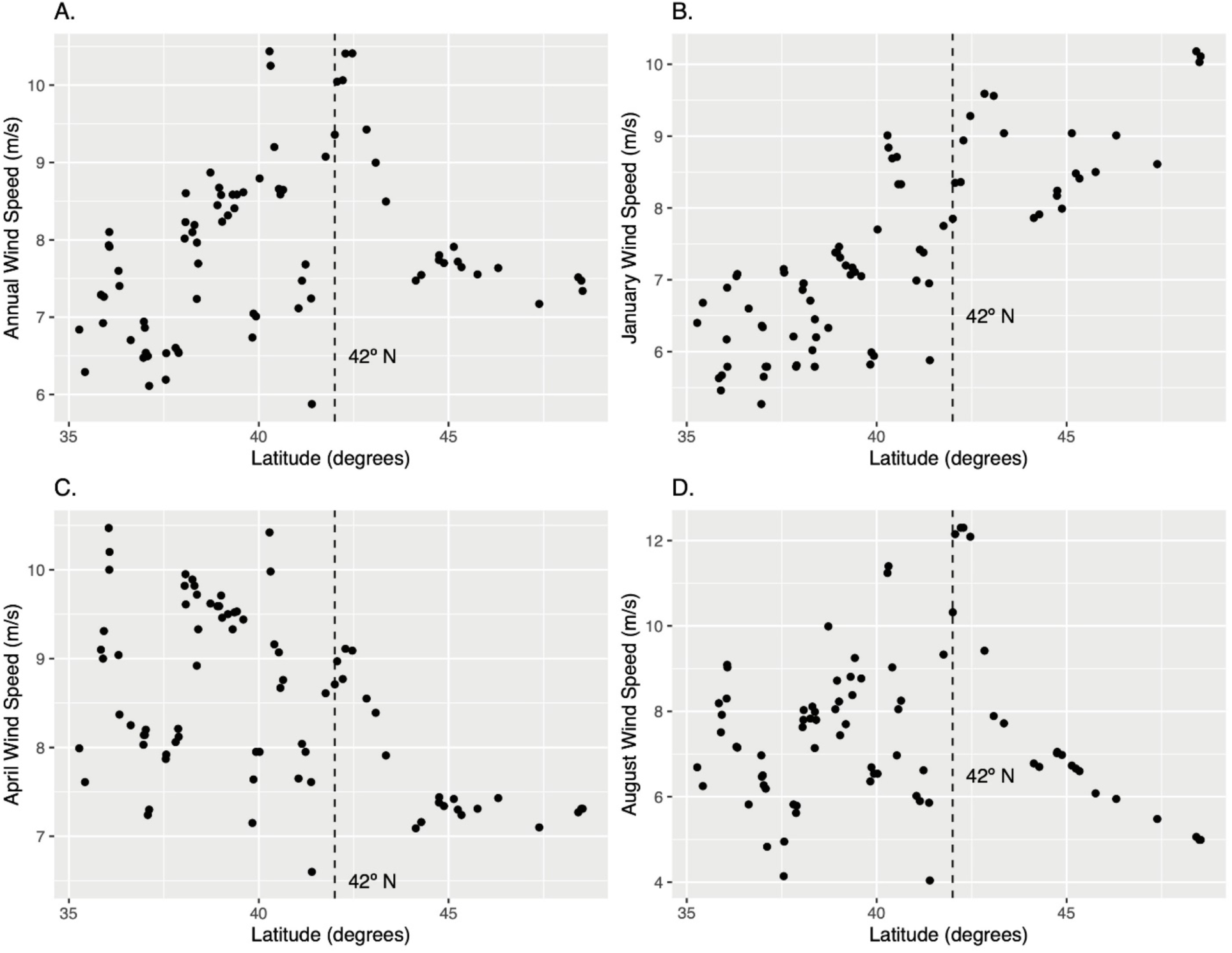
Mean wind speeds (m/s) along the United States Pacific Coast reported annually (A), during January (B), April (C), and August (D). The dashed line indicates 42° N, the latitude of the Oregon coast at which wind speeds start declining going further north.

However, the relationship between wind speeds and latitude varied dramatically based on time of year. Average wind speeds in Quarter 1 (*slope* = 0.19, *Multiple R²* = 0.46, *F*_1,76_ = 65.9, *P* < 0.001) and Quarter 4 (*slope* = 0.27, *Multipl*e *R²* = 0.50, *F*_1,76_ = 75.32, *P* < 0.001) of the year were strongly positively correlated with latitude. In contrast, Quarter 2 wind speeds (*slope* = -0.20, *Multiple R²* = 0.20, *F*_1,76_ = 19.1, *P* < 0.001) showed a negative relationship with latitude and Quarter 3 wind speeds were not correlated with latitude (*slope* = 0.05, *Multiple R²* = 0.01, *F*_1,76_ = 0.57, *P* > 0.05).

Finally, we constructed a linear model to evaluate how mean population height varied with annual mean wind speeds for each population from which accessions were collected. We also included the ocean exposure term in this model. While annual mean wind speed only had a marginally significant effect on plant height (*estimate* = -1.29, *P* = 0.073), open ocean exposure made a clear difference (*estimate* = -6.98, *P* < 0.001) and the model was a significant fit for the data (*Multiple R^2^* = 0.26, *F_2,62_* = 10.95, *P* < 0.001). In general, *M. guttatus* populations exposed to the open ocean were shorter than their protected counterparts (Fig. S1).

## DISCUSSION

In this study, we found a significant role of latitude and exposure to the open ocean in predicting genetically based trait variation among populations of the coastal perennial ecotype of *M. guttatus*. The trait most affected by these two factors was plant height, which was smaller for populations collected from locations exposed to the open ocean and was negatively correlated with latitude. Multiple environmental factors may have contributed to the evolution of these patterns of trait variation. These southern populations generally experience higher temperatures, less precipitation, and lower velocity winds than more northern populations, but this can vary based on the time of year. Further, populations exposed to the open ocean often experience greater wind speeds and more toxic salt spray, which is a byproduct of the wave energy of the ocean (Boyce 1954; Du & Hesp 2020). The results of this study further reinforce the idea that genetically-based phenotypic variation can be simultaneously partitioned in disjunct (among coastal perennial and inland annual ecotypes) and continuous, clinal ways (within ecotypes). We found similar overall results in a pilot experiment (See Appendix S1), indicating that our findings are replicable.

### Phenotypic variation is partitioned at multiple levels within species

Understanding the patterns by which genetic variation is partitioned among populations within a species across a geographic landscape has been a matter of considerable debate, especially in the classic plant evolutionary biological literature (reviewed in Lowry 2012). It is reasonable to hypothesize that the partitioning of that genetic variation is a function of how environmental variation is partitioned across the same landscape. However, several previous researchers have argued against the existence of ecotypes and that only clinal variation exists within species (Langlet 1971; Quinn 1978; Stebbins 1980; Levin 1993).

A series of prior studies has clearly established that the coastal populations of *Mimulus guttatus* from central California to southern Washington state collectively comprise a genetically distinct, geographically widespread ecotype (Lowry et al. 2008; Lowry and Willis 2010; Oneal et al. 2014; Twyford and Friedman 2015). This coastal perennial ecotype clusters as a distinct group from the rest of the *Mimulus guttatus* species complex based on both a set of morphological traits and allele frequencies of loci across the genome (Lowry 2012; Oneal et al. 2014; Twyford and Friedman 2015) but has not accumulated appreciable divergence through fixed genetic differences (Gould et al. 2017). This study demonstrates that while the coastal perennial ecotype is strikingly divergent from the inland annual ecotype, it also harbors genetically-based phenotypic variation that is partitioned along two major environmental gradients. Latitudinal clines within ecotypes/species have been observed in the classic literature for plant systems such as *Plantago maritima* (Gregor 1930, 1938) and *Achillea borealis* (Clausen et al. 1948).

Similarly, differences in traits including plant height have been found recently for coastal plants that occur on dunes versus headlands in *Senecio lautus* (Roda et al. 2013: James et al. 2021).

However, in contrast to *M. guttatus*, the evolution of short headland versus tall dune ecotypes in *S. latutus* has occurred independently through repeated parallel evolution (Roda et al. 2013; James et al. 2021) instead of occurring all within a single coastal ecotype. In the case of the coastal perennial ecotype of *M. guttatus*, the population genetic evidence suggests that it was derived at a single point in time from the rest of the species complex and then spread along the Pacific coast, adapting to local environments as it spread (Lowry et al. 2008; Twyford & Friedman 2015). During and following the spread of the coastal ecotype, there has likely been appreciable gene flow with other parts of the species complex, which is further supported by there only being four fixed nucleotide differences between these groups out of millions of SNPs (Gould et al. 2017).

### The environmental factors responsible for the geographic distribution of trait variation

While we have demonstrated that clines in plant height do occur within the coastal ecotype of *Mimulus guttatus*, the question remains as to what environmental factors have driven these clines. Though our analysis identified multiple factors correlated with both latitude and plant height, these factors must be carefully considered, as they are autocorrelated with each other and not all of them may have an impact on the selective advantage of plant height. Regardless of which environmental variables dictate the shape of the height cline, we have shown that exposure to the open ocean is predictive of shorter plant height in *M. guttatus* populations (Figs. 2, S1). These results are consistent with classic observations by Boyce (1954) and recent studies in *Setaria viridis* (Itoh 2021; Itoh et al. 2024).

Other researchers studying the evolution of ecotypes in the same region have found similar patterns in plant height variation along the Pacific coast of North America. These studies were far less comprehensive, but are useful, as they suggest that the patterns observed within the coastal ecotype of *M. guttatus* are not unique. For example, Clausen et al. (1948) found similar genetic variation with the coastal ecotype of *Achillea borealis* when comparing populations in Northern California, where populations north of the San Francisco Bay were shorter in stature than those from farther south in California in common garden experiments.

The causes of the latitudinal cline of plant height cannot be determined by this study alone and likely are complex. A global study of plant heights collected from an array of ecosystems found that there is a strong negative relationship between latitude and plant height (Moles et al. 2009).

However, none of the environmental variables evaluated in that study were very strongly correlated with plant height. For our intraspecific study, temperature and precipitation were the most highly correlated with latitude. Both of these variables could directly or indirectly be driving the evolution of plant height in this system. For example, the longer growing season in the southern end of the range of the coastal perennial ecotype could lead to greater competition, which would favor greater height. A positive relationship between temperature and plant height has been observed within species for altitudinal gradients (Totland & Birks 1996; Macek & Leps 2008; Pellissier et al. 2010), but other factors could also drive the overall negative association of elevation and height.

Perhaps the most compelling factor driving patterns of plant height is the combination of wind and salt spray. While wind is less closely correlated with latitude than other environmental factors, whether a population is exposed to the open ocean or is protected from salt spray by being located slightly inland can have a large effect on the impacts of wind and salt spray on plants (Barbour 1978; Du and Hesp 2020). Wind and salt spray have long been thought to drive the evolution of the stature of coastal plant species. For example, Boyce (1954) recognized that this compact growth form is advantageous for coastal plants because it helps them to avoid the impacts of salt spray by being more sheltered from the wind. Onshore wind from the ocean can be intense along the Pacific Coast, depending on the location and time of year (Grace 1977; Gilliam 2002). The intensity of wind speed increases with the distance above the ground (Grace 1977; Jaffe and Forbes 1993). While wind can be damaging by itself, it is particularly damaging to plants along ocean coastlines because it contains aerosolized oceanic salt spray (Boyce 1954; Barbour 1978; Cheplick and Demetri 1999; Griffiths et al. 2006; Kouali et al. 2017; Du and Hesp 2020). In winter months (Quarters 1 and 4, e.g. Fig 5b), there is a strong correlation between latitude and wind speeds, which could play a role in the evolution of shorter plant heights in more northern populations of this perennial plant ecotype. However, wind speed patterns are complex and their relationship with latitude shifts greatly over the year. Across the range of the coastal perennial ecotype, wind speeds and wave energy that drive salt spray are greater in the spring (April, May) than the summer (July, August) (Gilliam 2002; García-Reyes and Largier 2012). However, those winds speeds dissipate most during the summer in the region south of the San Francisco, especially in more protected areas, like the southern Monterey Bay (Fig. 3b), where plants in the field reach up to two meters in height (Lowry *personal observation*). In Northern California and Southern Oregon, high coastal windspeeds remain elevated throughout the summer months, which coincides with when this species flowers. Since reproductive structures are often the most sensitive to salt spray (Griffiths et al. 2006), maintaining a compact growth form when flowering under higher windspeed/salt spray conditions is likely to be advantageous. Field experimentation will be necessary to deconstruct how the time of year interacts with potential selection imposed by wind speeds and salt spray.

## CONCLUSIONS

Overall, our study demonstrates that there is dramatic and likely important phenotypic variation distributed within distinct ecotypes contained within species. Therefore, as in other species, like switchgrass (*Panicum virgatum*; Lowry et al. 2014), it is critical to account for both ecotypic and clinal variation when conceptualizing how functional genetic variation is distributed within a species. The exact causes of the clines within the coastal perennial ecotype of *M. guttatus* are still unclear, especially for trait correlations with latitude. However, the fact that exposure to the open ocean is predictive of plant height, suggests that wind and salt spray may play a crucial role in driving genetically-based variation in plant height in this system.

## ACKNOWLEDGEMENTS

This work is dedicated to the memory of Don Levin, whose papers and conversations challenged us to think hard about the nature of ecotypes. We would like to thank Mauricio Swartzendruber, Marty Schmidt, Simran Patel, Thomas Young, and Will Burke (Michigan State University) for assisting with plant care and data collection. We also want to thank all of the public agencies, private landowners, and First Nations Peoples who provided us with permission over the years to make collections for our research. Maddie James and two anonymous reviewers provided excellent comments on multiple versions of this manuscript, which greatly helped to improve it. Nate Emery, Anne-Sophie Bohrer, Linnea Fraser, Milagros del Pilar Jimenez-Hernandez, Jen Lau, Mackenzie Caple, and Emma Boehm provided advice with analysis using R. Caroline Draxl of NREL also shared her insight regarding the collection of offshore wind data. This work was funded by National Science Foundation Division of Integrative Organismal Systems Grants to DBL (IOS-1855927 and IOS-2153100).

## AUTHOR CONTRIBUTIONS

DBL conceived of and oversaw the experiments and analyses. TZ conducted the experiments and analyses. Both authors wrote the manuscript.

## DATA AVAILABILITY STATEMENT

All data will be made available at publication, either as supplementary information or on Dryad.

## SUPPORTING INFORMATION

Additional supporting information may be found online in the Supporting Information section at the end of the article.

Appendix S1: Pilot experiment and supplementary tables/figures

## Notes

### Competing Interest Statement

The authors have declared no competing interest.

### Summary of Updates

This manuscript has been extensively revised after two rounds of reviewer-driven revisions. The reviewers comments led us to think harder about our data and how best to approach analyses and interpretation. The most important breakthrough in rethinking our study was the realization that we could potentially test whether exposure of a population to the high wind speeds coming off of the open ocean versus being protected from that wind would have a predictive impact on plant height. Without looking at additional information, we divided all of the coastal perennial populations into exposed and protected populations. We then added this factor in our models and found that it was indeed highly predictive and improved the predictive capacity of latitude in the process. This new approach to our data has enhanced the value of our results and helped us to better interpret them in the discussion section of the manuscript. In addition to new approaches to our analyses, we have heavily revised the portion of the introduction and discussion of our manuscript that framed our research in terms of speciation. We have cut and rewritten much of these sections. Overall, we believe that our extensive reanalyses and revisions have greatly improved our manuscript and the value of this research.

## LITERATURE CITED

Barbour, M. G. 1978. Salt spray as a microenvironmental factor in the distribution of beach plants at Point Reyes, California. Oecologia 32: 213–224.

Barker, W. R., G. L. Nesom, P. M. Beardsle, and N. S. Fraga. 2012 A taxonomic conspectus of Phrymaceae: a narrowed circumscription for Mimulus, new and resurrected genera, and new names and combinations. Phytoneuron 39: 1–60.

Böcher, T. W. 1967. Continuous variation and taxonomy. Taxon 16: 255–258.

Boyce, S. G. 1954. The salt spray community. Ecological Monographs 24: 29–67.

Briggs, D., and S. M. Walters. 2016. Plant variation and evolution. Cambridge University Press, Cambridge, UK.

Cheplick, G. P., and H. Demetri. 1999. Impact of saltwater spray and sand deposition on the coastal annual *Triplasis purpurea* (Poaceae). American Journal of Botany 86: 703–710.

Clausen, J. 1951. Stages in the evolution of plant species. Cornell University Press, Ithaca, NY, USA.

Clausen, J., D. D. Keck, and W. M. Hiesey. 1948. Experimental studies on the nature of species. III. Environmental responses of climatic races of *Achillea*. Carnegie Institution of Washington, Washington, DC, USA.

Clausen, J., and W. M. Heisey. 1958. Experimental studies on the nature of species. IV. Genetic structure of ecological races. Carnegie Institution of Washington, Washington, DC, USA.

Draxl, C., B. M. Hodge, A. Clifton, and J. McCaa. 2015a. Overview and Meteorological Validation of the Wind Integration National Dataset Toolkit (Technical Report, NREL/TP-5000-61740). National Renewable Energy Laboratory, Golden, CO, USA.

Draxl, C., B. M. Hodge, A. Clifton, and J. McCaa. 2015b. The Wind Integration National Dataset (WIND) Toolkit. Applied Energy 151: 355–366.

Draxl, C., C. Philips, G. Scott, and W. Musial. 2017. “Characterization of Offshore Wind Resources in the Contiguous United States.” Presented at the 2017 NAWEA Conference, September 29, Ames, Iowa, USA. Retrieved July 18, 2022.

Du, J., and P. A. Hesp. 2020. Salt spray distribution and its impact on vegetation zonation on coastal dunes: a review. Estuaries and Coasts 43: 1885–1907.

Fick, S. E., and R. J. Hijmans. 2017. WorldClim 2: new 1km spatial resolution climate surfaces for global land areas. International Journal of Climatology 37: 4302–4315.

García-Reyes, M., and J. L. Largier. 2012. Seasonality of coastal upwelling off central and northern California: New insights, including temporal and spatial variability. Journal of Geophysical Research 117: C03028.

Gardiner, B., P. Berry, and B. Moulia. 2016. Review: Wind impacts on plant growth, mechanics and damage. Plant Science 245: 94–118.

Gilliam, H. 2002. Weather of the San Francisco Bay Region. University of California Press, Berkeley, CA, USA.

Gould, B. A., Y. Chen, and D. B. Lowry. 2017. Pooled ecotype sequencing reveals candidate genetic mechanisms for adaptive differentiation and reproductive isolation. Molecular Ecology 26: 163–177.

Grace, J. 1977. Plant Response to Wind. Academic Press, New York, NY, USA.

Gregor, J. W. 1930. Experiments on the genetics of wild populations. I. *Plantago maritima*. Journal of Genetics 22: 15–25.

Gregor, J. W. 1938. Experimental taxonomy II: Initial population differentiation in *Plantago maritima* L. of Britain. New Phytologist 37: 15–49.

Gregor, J. W., V. McM. Davey, and J. M. S. Lang. 1936. Experimental taxonomy I. Experimental garden technique in relation to the recognition of the small taxonomic units. New Phytologist 35: 323–350.

Griffiths, M. E., R. P. Keith, and C. M. Orians. 2006. Direct and indirect effects of salt spray and fire on coastal heathland plant physiology and community composition. Rhodora 108: 32–42.

Hall, M. C., C. J. Basten, and J. H. Willis. 2006. Pleiotropic quantitative trait loci contribute to population divergence in traits associated with life-history variation in *Mimulus guttatus*. Genetics 172: 1829–1844.

Hall, M. C., D. B. Lowry, and J. H. Willis. 2010. Is local adaptation in *Mimulus guttatus* caused by trade-offs at individual loci? Molecular Ecology 19: 2739–2753.

Heywood, V. H. 2011. The genesis of IOPB: a personal memoir. Taxon 60: 320–323.

Hitchcock, C. L., and A. Cronquist. 1973. Flora of the Pacific Northwest. University of Washington Press, Seattle, WA, USA.

Huxley, J. 1938. Clines: an auxiliary taxonomic principle. Nature 142: 219–220.

Itoh, M. 2021 Phenotypic variation and adaptation in morphology and salt spray tolerance in coastal and inland populations of *Setaria viridis* in central Japan. Weed Research 61:199–209.

Itoh M, Fukunaga K, Osako T. 2024. Local adaptation in parapatric and sympatric mosaic coastal habitats through trait divergence of *Setaria viridis*. Journal of Ecology 112: 784–799.

Jaffe, M. J., and S. Forbes. 1993 Thigmomorphogenesis: the effect of mechanical perturbation on plants. Plant Growth Regulation 12: 313–324.

James, M. E., M. J. Wilkinson, D. M. Bernal, H. Liu, H. L. North, J. Engelstädter, and D. Ortiz- Barrientos. 2021. Phenotypic and genotypic parallel evolution in parapatric ecotypes of Senecio. Evolution 75: 3115–3131.

Kaiser, H. F. 1960. The application of electronic computers to factor analysis. Educational and Psychological Measurement 20:141–151.

King, J., A. Clifton, and B. M. Hodge. 2014. Validation of Power Output for the WIND Toolkit (Technical Report, NREL/TP-5D00-61714). National Renewable Energy Laboratory, Golden, CO, USA.

Kouali, A., M. Mouradi, L. Latrach, A. Khadraji, O. Elasri, Z. Baicha, A. Berrichi, N. Kouddane, and A. Boukroute. 2017. Study of effects of seawater salt spray on growth, chlorophyll fluorescence and chlorophyll content in three coastal species of Morocco. Journal of Materials and Environmental Sciences 8: 4385–4390.

Langlet, O. 1963. Patterns and terms of intraspecific ecological variability. Nature 200: 347–348.

Langlet, O. 1971. Two hundred years of genecology. Taxon 20: 653–722.

Levin, D. A. 1993. Local speciation in plants: The rule not the exception. Systematic Botany 18: 197–208.

Lowry, D. B. 2010. Integrating genetics, geography, and local adaptation to understand ecotype formation in the yellow monkeyflower, *Mimulus guttatus*. PhD dissertation, Duke University, Durham, NC, USA.

Lowry, D. B. 2012. Ecotypes and the controversy over stages in the formation of new species. Biological Journal of the Linnean Society 106: 241–257.

Lowry, D. B., R. C. Rockwood, and J. H. Willis. 2008. Ecological reproductive isolation of coast and inland races of *Mimulus guttatus*. Evolution 62: 2196–2214.

Lowry, D. B., M. C. Hall, D. E. Salt, and J. H. Willis. 2009. Genetic and physiological basis of adaptive salt tolerance divergence between coastal and inland *Mimulus guttatus*. New Phytologist 183: 776–788.

Lowry, D. B., and J. H. Willis. 2010. A widespread chromosomal inversion polymorphism contributes to a major life-history transition, local adaptation, and reproductive isolation. PLoS Biology 8: e1000500.

Lowry, D. B, K. D. Behrman, P. Grabowski, G. P. Morris, J. R. Kiniry, T. E. Juenger. 2014. Adaptation between ecotypes and along environmental gradients in *Panicum virgatum*. The American Naturalist 183: 682–692.

Lowry, D. B., D. Popovic, D. J. Brennan, and L. M. Holeski. 2019. Mechanisms of a locally adaptive shift in allocation among growth, reproduction, and herbivore resistance in *Mimulus guttatus*. Evolution 73: 1168–1181.

Macek, P., and Leps, J. 2008. Environmental correlates of growth traits of the stoloniferous plant *Potentilla palustris*. Evolutionary Ecology 22: 419–435.

Nesom, G. L. 2012. Taxonomy of *Erythranthe* sect. Simiola (Phrymaceae) in the USA and Mexico. Phytoneuron 2012–40: 1–123.

Nesom, G. L. 2014. Further observations on relationships in the *Erythranthe guttata* group (Phrymaceae). Phytoneuron 2014–93: 1–8

Oneal, E., D. B. Lowry, K. M. Wright, Z. Zhu, and J. H. Willis. 2014. Divergent population structure and climate associations of a chromosomal inversion polymorphism across the *Mimulus guttatus* species complex. Molecular Ecology 23: 2844–2860.

Pellissier, L., B. Fournier, A. Guisan, and P. Vittoz P. 2010. Plant traits co-vary with altitude in grasslands and forests in the European Alps. Plant Ecology 211: 351–365.

Posit Team. 2023. RStudio: Integrated Development Environment for R. Posit Software, PBC, Boston, MA. URL http://www.posit.co/.

Quinn, J. A. 1978 Plant ecotypes: ecological or evolutionary units? Bulletin of the Torrey Botanical Club 1: 58–64.

R Core Team. 2022. R: A language and environment for statistical computing. R Foundation for Statistical Computing, Vienna, Austria. URL https://www.R-project.org/.

Roda, F., L. Ambrose, G. M. Walter, H. L. Liu, A. Schaul, A. Lowe, P. B. Pelser, P. Prentis, L. H. Rieseberg, and D. Ortiz-Barrientos. 2013. Genomic evidence for the parallel evolution of coastal forms in the *Senecio lautus* complex. Molecular Ecology 22: 2941–2952.

Stebbins, G. L. 1980. Botany and the synthetic theory of evolution. p139-152. In: Mayr E Provine WB, eds. Evolutionary synthesis: perspectives on the unification of biology. Harvard University Press, Cambridge, MA, USA.

Telewski, F. W. 2021. Mechanosensing and plant growth regulators elicited during the thigmomorphogenetic response. Frontiers in Forests and Global Change 3: 574096.

Totland, O., and Birks, H.J.B. 1996. Factors influencing inter-population variation in *Ranunculus acris* seed production in an alpine area of southwestern Norway. Ecography 19: 269–278.

Turesson, G. 1922a. The species and the variety as ecological units. Hereditas 3: 100–113.

Turesson, G. 1922b. The genotypic response of the plant species to habitat. Hereditas 3: 211–350.

Twyford, A. D., and J. Friedman. 2015. Adaptive divergence in the monkey flower *Mimulus guttatus* is maintained by a chromosomal inversion. Evolution 69: 1476–1486.

Vickery, R. K. 1952. A study of the genetic relationships in a sample of the *Mimulus guttatus* complex. PhD Dissertation, Stanford University, Stanford, CA, USA.

Vickery, R. K. 1978. Case studies in the evolution of species complexes in *Mimulus*. Evolutionary Biology 11: 405–507.

Wu, C. A., D. B. Lowry, A. M. Cooley, K. M. Wright, Y. W. Lee, and J. H. Willis. 2008. *Mimulus* is an emerging model system for the integration of ecological and genomic studies. Heredity 100: 220–230.

