## Appendix for "Multiple clines within an ecotype of the yellow monkeyflower, *Mimulus guttatus*"

### APPENDIX S1: Pilot experiment and supplementary tables/figures

***Pilot experiment methods –***

We initially conducted a pilot experiment with fewer plants to begin our investigation of potential clines in *M. guttatus* phenotypes. We planted seeds collected from coastal perennial *M. guttatus* according to maternal family ($N$ = 135 individuals, 135 seed families, 56 populations) in 3.5 sq. in. pots with SUREMIX (Michigan Grower Products, Inc., Galesburg, Michigan, USA). We included a variable number of seed families per population for the pilot experiment as well (Range = 1-4, Mean = 2.41 families); each seed family was represented by one individual. We randomized these pots into flats (17 pots per flat) and stratified the seeds in a dark cold room (set to 4º C) for a week to encourage more uniform germination before being transferred to the greenhouses at Michigan State University. All germinating seedlings were removed from each pot, except for the centermost plant. Plants were bottom-watered approximately every other day throughout the remainder of the experiment. Plants were grown in the greenhouses with 16 h of supplemental lighting and room temperature set to 22º C. We recorded approximate germination dates and flowering dates, and measured plant flowering height (cm) and stem width (mm) for each plant at the time of first flower.

We also gathered climate data for each population via the WorldClim database (Fick and Hijmans 2017) and approximated offshore wind velocity at each population's latitude using the NREL Wind Prospector (King et al. 2014; Draxl et al. 2015a,b). Details of these procedures can be found in the main Methods section of this study.

To assess the relationship of plant traits with population latitude and environmental variables, we conducted linear regression and PCA analyses as described in the main Methods section of this study. Our analyses of the pilot project data also accounted for whether populations were exposed to or protected from the open ocean by geographical features by using a binary factor in our models. All analyses were performed with R statistical software (v.4.2.1, R Core Team 2022).

***Pilot experiment results –***

In our pilot experiment, we found that there was a significant negative relationship between latitude of origin and mean height in coastal perennial *M. guttatus* populations (*Multiple* $R^2$ = 0.29, $F_{2,53}$ = 11.06, $P$ < 0.001; Fig. S2). Both latitude and ocean exposure had significant effects on mean population height (Table S3). We also found a significant model fit predicting mean population stem width from latitude (*Multiple* $R^2$ = 0.24, $F_{2,52}$ = 8.09, $P$ < 0.001), but the exposure term was not significant (Table S3). Despite including a significant term for latitude as a predictor of flowering time, that model only had a marginally significant fit (*Multiple* $R^2$ = 0.09, $F_{2,53}$ = 2.66, $P$ = 0.079; Table S3). The relationship between latitude and plant height was the strongest and so was the focus of our subsequent analyses.

To gain further insight into the possible causes of the negative relationship between latitude and plant height, we conducted principal component analyses of climate factors. For our pilot experiment, we focused our analyses on climate PC1 through PC4, as all of them had eigenvalues greater than 1. These four PCs accounted for 94.55% of cumulative variance in the climate factors; PC1 explained 59.03% of the variation, followed by PC2 at 19.57%, PC3 at

11.13%, and PC4 at 4.82%. We found that PC1 had a significant negative relationship with plant height (*Multiple* $R^2 = 0.28$, $F_{2,53} = 10.47$, $P < 0.001$), with latitude and ocean exposure both being significant predictors in the model (Table S4). We assessed nonlinear models for PC1 due to the nonlinear relationship of wind speed and latitude but confirmed that the linear model fit the data best (*AIC linear* = 343.80, *AIC quadratic* = 345.73, *AIC cubic* = 345.35). Models including the other three PCs were significant fits for the data, but none of these included PC as a significant predictor (Table S4). We also compared the model with just PC1 to a model including all four PCs as predictors, but again found the PC1 model to be the best fit for the data via AIC (*AIC PC1* = 343.80, *AIC All PCs* = 347.14). We focused the remainder of our initial analyses on PC1, as it was the only model to include its principal component as a significant predictor and was the best model fit per AIC.

Given that PC1 had a significant negative correlation with height, we investigated which climate variables were most highly loaded on this PC axis. The most heavily loaded variables on PC1 were Annual Precipitation, Average Annual Temperature, Precipitation of the Wettest Quarter, Precipitation of the Coldest Quarter, Precipitation of the Driest Quarter, Temperature of the Coldest Quarter, Precipitation Seasonality, Precipitation of the Warmest Quarter, and Precipitation of the Wettest Month. Annual Temperature was negatively correlated with PC1, while all others were positively correlated. While not as heavily loaded, Average Wind Speeds in Quarter 1 (January, February, March) and Quarter 4 (October, November, December) were both positively correlated with PC1. Overall, a higher value of PC1 indicates that populations at those locations experience more precipitation, greater wind speeds, and lower temperatures. More specific information regarding loadings of individual climate variables on PC1 can be found in Table S5.

Using the principal components from our pilot experiment, we found that PC1 had a very strong positive relationship with latitude (*slope* = 0.76, *Multiple* $R^2 = 0.93$, $F_{1,54} = 689$, $P < 0.001$). The other PCs were not significantly correlated with latitude ($P > 0.05$). We also regressed climate factors that were highly-loaded on PC1 against latitude to determine how well variation in each individual climate factor was predicted by latitude (Table S6).

To explore our hypothesized effect of wind speed on mean population heights, we constructed a linear model and included a binary factor for exposure to the open ocean. While mean annual wind speed was not a significant term in our model predicting mean population heights ($F_{1,53} = 1.18$, $P > 0.05$), *M. guttatus* exposed to the ocean were significantly shorter (*estimate* = -4.35, $F_{1,53} = 8.74$, $P = 0.005$; Fig. S3). Overall, the model was a significant fit for the data (*Multiple* $R^2 = 0.15$, $F_{2,53} = 4.84$, $P = 0.012$).

***Supplemental tables and figures –***

**Table S1.** *M. guttatus* population codes, coordinates, and experiments in which they were used

| PopCode | Latitude | Longitude | Experiment |
|---|---|---|---|
| AGG | 52.62567 | -131.4218 | Main |
| ALA | 58.00433 | -135.7425 | Main |
| BCB | 36.06285 | -121.5922 | Main |
| BHE | 38.30703 | -123.0584 | Both |
| BLN | 42.83772 | -124.5601 | Both |
| BND | 43.07971 | -124.43 | Both |
| BOB | 48.52853 | -124.451 | Main |
| BON | 37.00016 | -122.182 | Both |
| BRK | 42.06429 | -124.3011 | Both |
| BYR | 43.34902 | -124.3458 | Both |
| CAB | 39.3592 | -123.8168 | Both |
| CKI | 45.24213 | -123.9687 | Main |
| CMD | 40.40923 | -124.392 | Both |
| CML | 41.04636 | -124.1226 | Both |
| COV | 40.02157 | -124.0688 | Both |
| CPB | 53.17103 | -131.7848 | Main |
| CPT | 36.9746 | -121.9451 | Both |
| CRD | 36.30686 | -121.8503 | Both |
| CRU | 35.84372 | -121.4039 | Both |
| CVG | 38.3723 | -123.0552 | Both |
| CVR | 38.3684 | -123.0606 | Pilot |
| DAV | 37.02498 | -122.2175 | Both |
| DIL | 38.25167 | -122.9678 | Main |
| DIS | 46.3053 | -124.072 | Both |
| FPT | 37.80918 | -122.4751 | Both |
| FRA | 39.0083 | -123.694 | Main |
| GBM | 41.37863 | -124.0696 | Both |
| GRE | 37.07795 | -122.2677 | Both |
| GRZ | 40.56885 | -124.2118 | Both |
| GWC | 37.56394 | -122.5138 | Both |
| HEC | 44.13536 | -124.1235 | Both |
| HOC | 47.3854 | -123.1473 | Main |
| HSI | 52.58167 | -131.445 | Main |
| HUM | 39.92726 | -123.7608 | Main |
| KUN | 52.75667 | -131.5694 | Main |
| LAG | 41.23157 | -124.1064 | Both |
| LBG | 49.07185 | -125.7676 | Main |
| LJA | 39.86053 | -123.9024 | Both |
| LOK | 45.33825 | -123.9771 | Both |
| MTO | 40.28343 | -124.3596 | Both |
| MTR | 37.5517 | -122.5137 | Both |
| MUR | 37.87685 | -122.5578 | Pilot |
| MXX | 40.31068 | -124.2671 | Both |
| NAV | 39.1869 | -123.7572 | Main |
| OPB | 42.46402 | -124.4229 | Both |
| ORO | 35.27335 | -120.8891 | Both |
| OSW | 45.76108 | -123.9666 | Both |
| OTT | 44.75224 | -124.0641 | Main |
| OZF | 38.95578 | -123.6603 | Pilot |
| PCR | 41.39856 | -124.0688 | Main |
| PGR | 36.62799 | -121.9208 | Both |
| PPT | 41.13709 | -124.1519 | Both |
| PTL | 42.28225 | -124.405 | Both |
| PTR | 38.04861 | -122.8694 | Both |
| PTU | 44.27916 | -124.1121 | Both |
| PUN | 44.74484 | -124.06 | Both |
| REI | 52.88056 | -131.5167 | Main |
| SCC | 48.41262 | -124.0121 | Main |
| SDB | 35.92127 | -121.4668 | Both |
| SIM | 44.87659 | -124.0383 | Both |
| SLI | 49.89612 | -124.5907 | Main |
| SRN | 38.72593 | -123.4805 | Both |
| STB | 37.89132 | -122.6347 | Both |
| SUR | 36.33194 | -121.8887 | Both |
| SWB | 39.03598 | -123.6905 | Main |
| TIT | 52.785 | -131.5836 | Main |
| TSU | 41.75983 | -124.2221 | Both |
| USB | 39.83218 | -123.8493 | Both |
| VIC | 36.04834 | -121.5885 | Both |
| WCK | 42.00256 | -124.2129 | Both |
| WCR | 35.89838 | -121.4613 | Both |
| WEV | 35.42313 | -120.7887 | Both |
| WOD | 39.31239 | -123.8029 | Both |
| WPT | 42.21213 | -124.3767 | Both |
| WPV | 39.59383 | -123.7854 | Both |
| WSZ | 36.96233 | -122.0764 | Both |
| WTB | 38.40525 | -123.0961 | Both |

Table S2. Principal component loadings and coordinates (main experiment)

| Climate Variables | PC1 loading | PC2 loading | PC3 loading | PC1 coordinate | PC2 coordinate | PC3 coordinate |
|---|---|---|---|---|---|---|
| Q1Wind | 4.446 | 3.064 | 1.878 | 0.776 | 0.41 | -0.221 |
| Q2Wind | 0.538 | 4.073 | 23.261 | -0.27 | 0.472 | -0.776 |
| Q3Wind | 0.61 | 6.308 | 18.032 | 0.287 | 0.588 | -0.683 |
| Q4Wind | 4.969 | 3.552 | 1.422 | 0.82 | 0.441 | -0.192 |
| WindAnnual | 1.351 | 6.904 | 16.059 | 0.428 | 0.615 | -0.645 |
| AnnTemp | 6.718 | 0.166 | 0.005 | -0.953 | 0.095 | -0.011 |
| DiurnalRange | 3.847 | 3.591 | 5.202 | -0.722 | -0.443 | -0.367 |
| Isothermality | 4.57 | 3.211 | 0.454 | -0.786 | 0.419 | -0.108 |
| TempSeasonality | 1.18 | 13.5 | 2.592 | 0.4 | -0.86 | -0.259 |
| MaxTemp_WarmMonth | 2.991 | 8.281 | 4.862 | -0.636 | -0.673 | -0.355 |
| MinTemp_ColdMonth | 1.862 | 9.333 | 1.688 | -0.502 | 0.715 | 0.209 |
| TempAnnRange | 0.633 | 13.563 | 5.572 | -0.293 | -0.862 | -0.38 |
| TempWetQuarter | 5.01 | 4.892 | 0.142 | -0.823 | 0.517 | 0.061 |
| TempDryQuarter | 3.051 | 6.88 | 3.184 | -0.643 | -0.614 | -0.287 |
| TempWarmQuarter | 3.659 | 5.741 | 2.214 | -0.704 | -0.561 | -0.239 |
| TempColdQuarter | 5.26 | 4.118 | 0.308 | -0.844 | 0.475 | 0.089 |
| AnnPrecip | 6.661 | 0.06 | 0.951 | 0.949 | -0.057 | -0.157 |
| PrecipWetMonth | 6.018 | 0.048 | 2.143 | 0.902 | -0.052 | -0.236 |
| PrecipDryMonth | 5.231 | 1.434 | 2.82 | 0.841 | -0.28 | 0.27 |
| PrecipSeasonality | 6.481 | 0.128 | 0.648 | -0.937 | 0.084 | -0.13 |
| PrecipWetQuarter | 6.264 | 0.054 | 1.891 | 0.921 | -0.054 | -0.221 |
| PrecipDryQuarter | 6.151 | 0.554 | 1.349 | 0.912 | -0.174 | 0.187 |
| PrecipWarmQuarter | 6.425 | 0.499 | 0.941 | 0.932 | -0.165 | 0.156 |
| PrecipColdQuarter | 6.074 | 0.045 | 2.382 | 0.907 | -0.05 | -0.248 |

**Table S3.** ANOVA and linear regression summaries for *M. guttatus* traits in the pilot greenhouse. Asterisks denote significance at p<0.05 (*), p<0.01 (**), and p<0.001 (***).

| Trait | Parameter Summaries | | | | | | Full Model Summaries | | | |
|---|---|---|---|---|---|---|---|---|---|---|
| | Predictor | Estimate | Sum Sq. | df | F | P | Multiple $R^2$ | df | F | P |
| Flowering Time (days) | Latitude | 0.4488 | 89.1 | 1,53 | 4.650 | 0.0356* | 0.0913 | 2,53 | 2.661 | 0.079 |
| | Exposure | 0.2221 | 0.62 | 1,53 | 0.032 | 0.8585 | | | | |
| Stem Width (mm) | Latitude | -0.1352 | 7.74 | 1,52 | 13.716 | 5.16e-04*** | 0.2372 | 2,52 | 8.087 | 8.749e-04*** |
| | Exposure | -0.0711 | 0.06 | 1,52 | 0.107 | 0.7455 | | | | |
| Height (cm) | Latitude | -0.8128 | 292.17 | 1,53 | 11.934 | 0.0011** | 0.2944 | 2,53 | 11.06 | 9.702e-05*** |
| | Exposure | -2.9247 | 106.73 | 1,53 | 4.359 | 0.0416* | | | | |

**Table S4.** ANOVA, linear regression, and AIC summaries for models evaluating the relationship of selected principal components to *M. guttatus* height in the pilot greenhouse. Asterisks denote significance at p<0.5 (*), p<0.01 (**), and p<0.001 (***). Lowest AIC is bolded.

| Principal Component | Parameter Summaries | | | | | | Full Model Summaries | | | | AIC |
|---|---|---|---|---|---|---|---|---|---|---|---|
| | Predictor | Estimate | Sum Sq. | df | F | P | Multiple $R^2$ | df | F | P | |
| PC1 | Dim. 1 | -0.6109 | 271.73 | 1,53 | 10.927 | 0.002** | 0.2833 | 2,53 | 10.47 | 1.468e-04*** | **343.799** |
| | Exposure | -3.1012 | 121.79 | 1,53 | 4.898 | 0.031* | | | | | |
| PC2 | Dim. 2 | 0.9522 | 77.4 | 1,52 | 2.736 | 0.104 | 0.1998 | 3,52 | 4.327 | 0.008** | 351.970 |
| | Exposure | -4.4986 | 268 | 1,52 | 9.469 | 0.003** | | | | | |
| | Interaction | -1.4443 | 118.2 | 1,52 | 4.176 | 0.046* | | | | | |
| PC3 | Dim. 3 | 0.487 | 33.23 | 1,53 | 1.131 | 0.292 | 0.1536 | 2,53 | 4.809 | 0.012* | 353.113 |
| | Exposure | -3.9421 | 200.82 | 1,53 | 6.838 | 0.012* | | | | | |
| PC4 | $(\text{Dim. }4)^2$ | 12.043 | 161.57 | 2,52 | 2.941 | 0.062 | 0.2234 | 3,52 | 4.986 | 0.004** | 350.294 |
| | Exposure | -4.135 | 230.35 | 1,52 | 8.387 | 0.006** | | | | | |

**Table S5.** Principal component loadings and coordinates (pilot experiment)

| Climate Variables | PC1 loading | PC2 loading | PC3 loading | PC4 loading | PC1 coordinate | PC2 coordinate | PC3 coordinate | PC4 coordinate |
|---|---|---|---|---|---|---|---|---|
| Q1Wind | 4.745 | 3.893 | 1.893 | 0.007 | 0.82 | -0.428 | 0.225 | 0.009 |
| Q2Wind | 0.012 | 2.341 | 29.701 | 0.048 | -0.042 | -0.332 | 0.891 | 0.024 |
| Q3Wind | 1.774 | 3.333 | 17.748 | 0.01 | 0.501 | -0.396 | 0.689 | -0.011 |
| Q4Wind | 5.28 | 3.177 | 1.023 | 0.021 | 0.865 | -0.386 | 0.165 | -0.015 |
| WindAnnual | 2.591 | 4.445 | 15.244 | 0 | 0.606 | -0.457 | 0.638 | 0 |
| AnnTemp | 6.343 | 0.22 | 0.011 | 4.128 | -0.948 | -0.102 | -0.017 | -0.218 |
| DiurnalRange | 3.497 | 4.888 | 5.302 | 9.528 | -0.704 | 0.479 | 0.376 | 0.332 |
| Isothermality | 3.898 | 2.213 | 1.848 | 18.127 | -0.743 | -0.322 | 0.222 | 0.458 |
| TempSeasonality | 0.463 | 17.639 | 1.789 | 3.794 | 0.256 | 0.91 | 0.219 | -0.209 |
| MaxTemp_WarmMonth | 3.926 | 7.093 | 3.338 | 0.355 | -0.746 | 0.577 | 0.299 | -0.064 |
| MinTemp_ColdMonth | 1.178 | 11.607 | 2.268 | 17.63 | -0.409 | -0.738 | -0.246 | -0.452 |
| TempAnnRange | 1.453 | 13.62 | 4.744 | 1.651 | -0.454 | 0.8 | 0.356 | 0.138 |
| TempWetQuarter | 4.751 | 5.731 | 0.096 | 1.339 | -0.82 | -0.519 | -0.051 | -0.124 |
| TempDryQuarter | 3.991 | 4.896 | 1.199 | 11.223 | -0.752 | 0.48 | 0.179 | -0.36 |
| TempWarmQuarter | 4.441 | 3.987 | 0.643 | 10.368 | -0.793 | 0.433 | 0.131 | -0.346 |
| TempColdQuarter | 5.005 | 4.811 | 0.266 | 0.633 | -0.842 | -0.475 | -0.084 | -0.086 |
| AnnPrecip | 6.536 | 0.439 | 0.375 | 2.297 | 0.962 | 0.144 | 0.1 | -0.163 |
| PrecipWetMonth | 5.966 | 0.425 | 1.1 | 5.906 | 0.919 | 0.141 | 0.171 | -0.261 |
| PrecipDryMonth | 4.448 | 2.012 | 4.369 | 1.957 | 0.794 | 0.307 | -0.342 | 0.15 |
| PrecipSeasonality | 5.964 | 0.319 | 1.005 | 1.231 | -0.919 | -0.122 | 0.164 | -0.119 |
| PrecipWetQuarter | 6.228 | 0.423 | 0.946 | 4.013 | 0.939 | 0.141 | 0.159 | -0.215 |
| PrecipDryQuarter | 5.561 | 1.154 | 2.309 | 0.467 | 0.888 | 0.233 | -0.248 | 0.073 |
| PrecipWarmQuarter | 5.863 | 0.904 | 1.535 | 0.471 | 0.911 | 0.206 | -0.203 | 0.074 |
| PrecipColdQuarter | 6.085 | 0.431 | 1.248 | 4.795 | 0.928 | 0.142 | 0.183 | -0.235 |

**Table S6.** Regressions of climate factors that were highly-loaded on PC1 against latitude (pilot experiment)

| *Climate Factors* | Estimate | F | Multiple $R^2$ | P |
|---|---|---|---|---|
| Annual Precipitation | 0.0459 | 401.9 | 0.8816 | < 2.2e-16 |
| Coldest Quarter Precipitation | 0.1038 | 200.9 | 0.7882 | < 2.2e-16 |
| Driest Quarter Precipitation | 0.5854 | 436.9 | 0.89 | < 2.2e-16 |
| Mean Annual Temperature | -2.3367 | 237.2 | 0.8146 | < 2.2e-16 |
| Mean Temperature, Coldest Quarter | -1.8891 | 163.2 | 0.7514 | < 2.2e-16 |
| Mean Wind Speed, 1st Quarter | 2.1737 | 50.06 | 0.4811 | 3.11E-09 |
| Mean Wind Speed, 4th Quarter | 1.6486 | 74.6 | 0.5801 | 9.37E-12 |
| Precipitation Seasonality | -0.2794 | 607.2 | 0.9183 | < 2.2e-16 |
| Warmest Quarter Precipitation | 0.5861 | 470.4 | 0.897 | < 2.2e-16 |
| Wettest Month Precipitation | 0.2877 | 191.4 | 0.7799 | < 2.2e-16 |
| Wettest Quarter Precipitation | 0.0978 | 239.6 | 0.8161 | < 2.2e-16 |

Note: F-statistic degrees of freedom were all 1,54

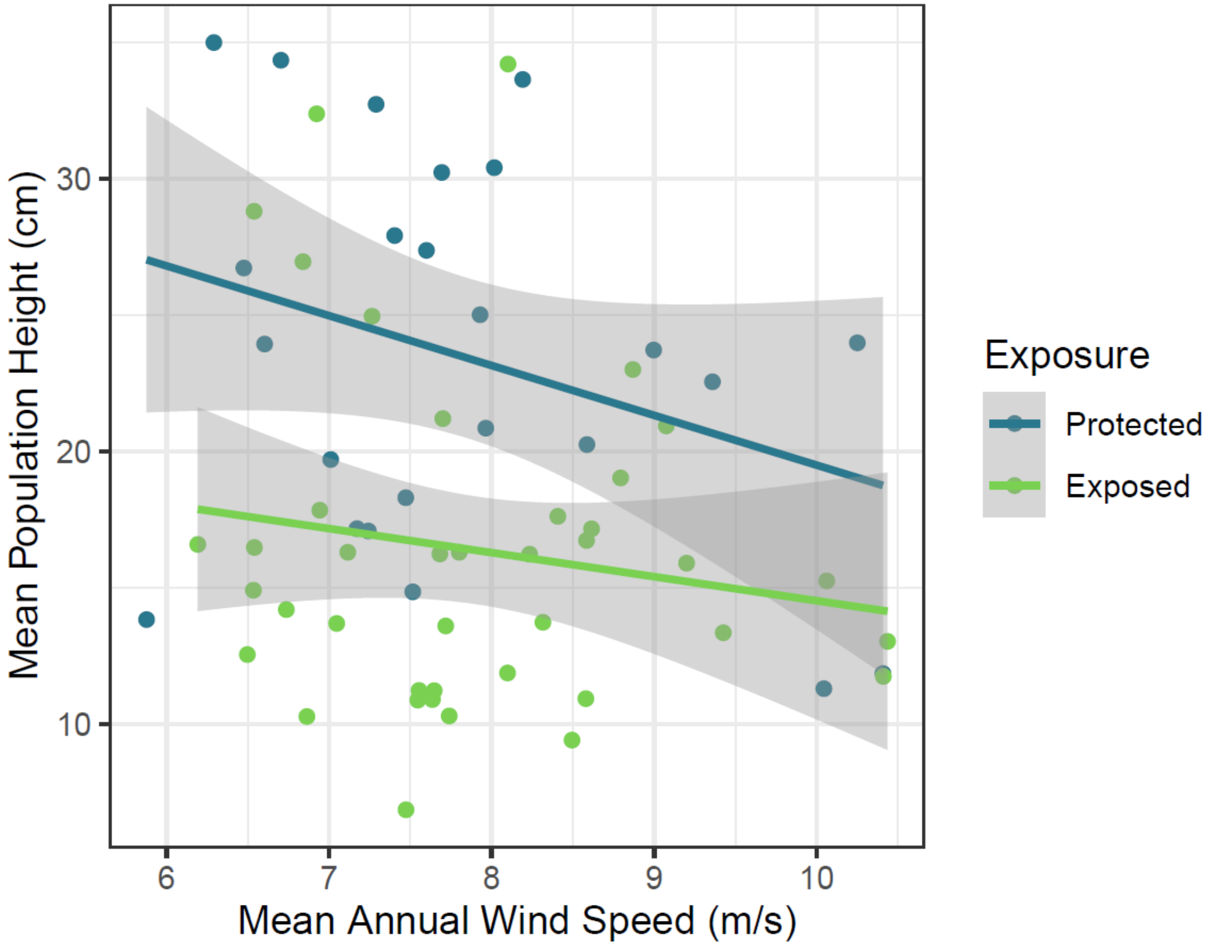


**Figure S1.** Annual wind speed had a marginally negative effect on mean *M. guttatus* population height in the main experiment ($F_{1,62} = 3.32$, $P = 0.073$), but populations exposed to the open ocean were significantly shorter than those protected from the wind ($F_{1,62} = 18.19$, $P < 0.001$). The overall model demonstrated a significant fit for our data ($R^2 = 0.26$, $F_{2,62} = 10.95$, $P < 0.001$).

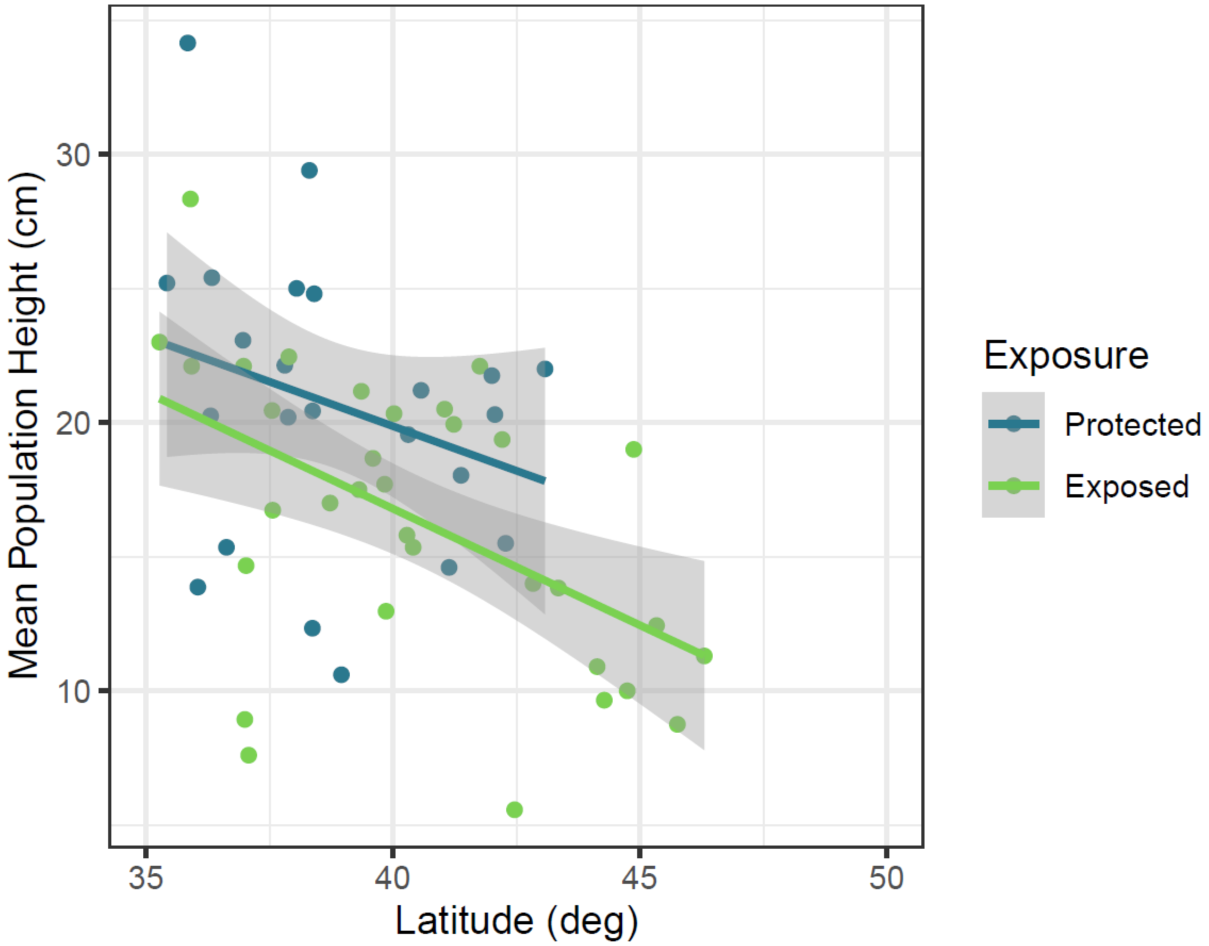


**Figure S2.** Mean *M. guttatus* population height decreased significantly with latitude of origin in the pilot experiment ($P = 0.001$); populations exposed to the ocean were shorter than protected populations ($P = 0.042$), but both maintain the same relationship between latitude and height. The relationship between latitude (in degrees) and mean height in *M. guttatus* populations was determined by linear regression.

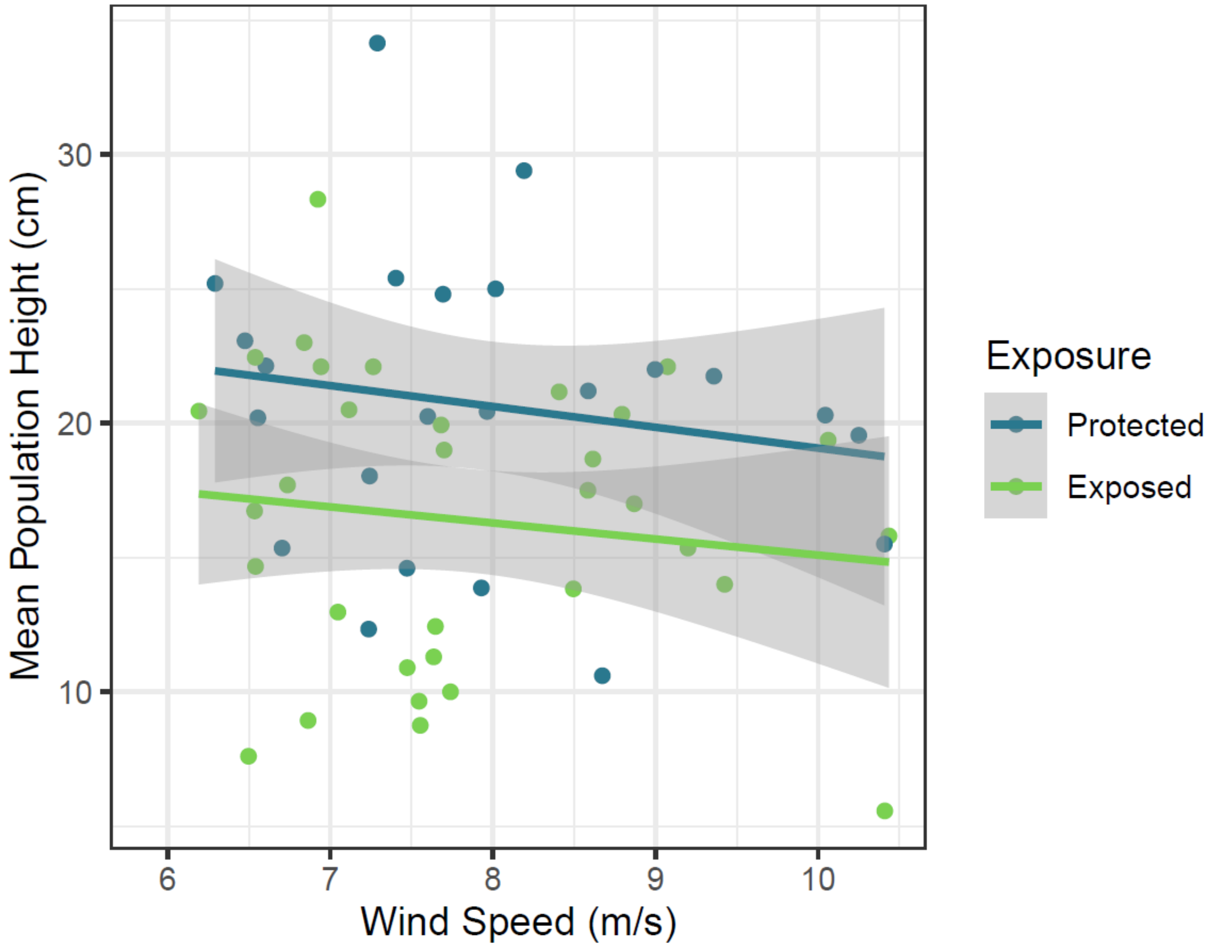


**Figure S3.** Annual mean wind speed did not have a significant effect on *M. guttatus* mean population height in the pilot experiment ($F_{1,53}$ = 1.18, $P$ = 0.282), but populations exposed to the open ocean were notably shorter ($F_{1,53}$ = 8.74, $P$ = 0.005). The overall model demonstrated a significant fit to our data ($R^2$ = 0.15, $F_{2,53}$ = 4.84, $P$ = 0.012).
